# Benchmarking Relational Graph Convolutional Networks for Glioblastoma Biomarker Discovery via ceRNA and Copy Number Variation Analysis

**DOI:** 10.64898/2026.08.16.744525

**Authors:** Samarth Khandelwal, Justin Zhan, Nicholas Jarvis

## Abstract

Glioblastoma (GBM) is a highly aggressive brain tumor with a five-year survival rate of 6.9%, attributable in substantial part to the shortage of reliable biomarkers. Competing endogenous RNA (ceRNA) and copy number variation (CNV) analyses each carry biomarker-identification potential, but existing work treats them separately. We built a heterogeneous ceRNA graph over 2,075 differentially expressed loci, added gene-level copy number, and evaluated relational graph convolutional networks (RGCNs) against graph and non-graph baselines under 10-fold cross-validation. The RGCN discriminated best among the graph architectures tested (AUCROC 0.874 *±* 0.070), and a late-fusion ensemble with the CNV branch scored highest overall (0.883 *±* 0.072). We then introduce a set of controls that separate network signal from properties of the label set. Positive labels are concentrated on miRNA nodes (26 of 31; 8.28% of miRNA nodes against 0.26% of mRNA nodes), and node type is recoverable from the relation structure. A logistic regression on one-hot node type alone, using no topology, reaches 0.824 *±* 0.139. An RGCN retrained on labels permuted within node type matches the observed PR-AUC (*p* = 0.33) but not the observed AUCROC (0.799 against 0.876, *p* = 0.015), and within the miRNA subgraph discrimination is weak and not distinguishable from that null (*p* = 0.10). The fusion gain has the same source: copy number is undefined at all 314 miRNA loci, the `cnv_observed` flag alone reaches 0.697, and the copy-number value with that flag removed reaches 0.475 *±* 0.007. Under this label set and graph construction, then, most of the apparent performance is carried by a node-type prior, with a modest residual network signal on AUCROC. The controls are inexpensive, require no additional data, and apply to any graph model trained on curated labels; we release the graph, labels, and controls so that they can be run in related designs. The five candidate loci remain differentially expressed when re-estimated from raw read counts and are supported by an alias-expanded literature screen, both independent of the model; their prognostic value is not assessed here.

## 1 INTRODUCTION

Brain and central nervous system cancers caused an estimated 248,305 deaths worldwide in 2022 (Bray et al., 2024). Among them, glioblastoma is the most lethal and common malignant form, accounting for over half of malignant primary brain tumors, with a 6.9% five-year survival rate (Ostrom et al., 2023). Much of that outcome traces to the lack of reliable biomarkers, which blocks accurate patient subtyping and diagnosis and leaves clinicians applying the Stupp protocol uniformly, regardless of tumor responsiveness (Roncevic et al., 2025).

Identifying biomarkers in glioblastoma remains a challenge, primarily due to the complex nature of gene regulation. Current statistical approaches can miss these molecular interactions, and few clinically useful targets have emerged. Better frameworks for identifying glioblastoma biomarkers are therefore needed.

### 1.1 Competing Endogenous RNA (ceRNA) Mechanisms in Glioblastoma

Noncoding RNAs cross-regulate one another through shared microRNA binding sites, forming what is known as the ceRNA network (Salmena et al., 2011). A microRNA (miRNA) bound to a messenger RNA (mRNA) silences it and prevents its translation into protein. Long non-coding RNA (lncRNA) and other transcripts carrying the same binding sites compete for that miRNA, and in sponging it they release the mRNA for translation (Figure 1).

**Figure 1.**
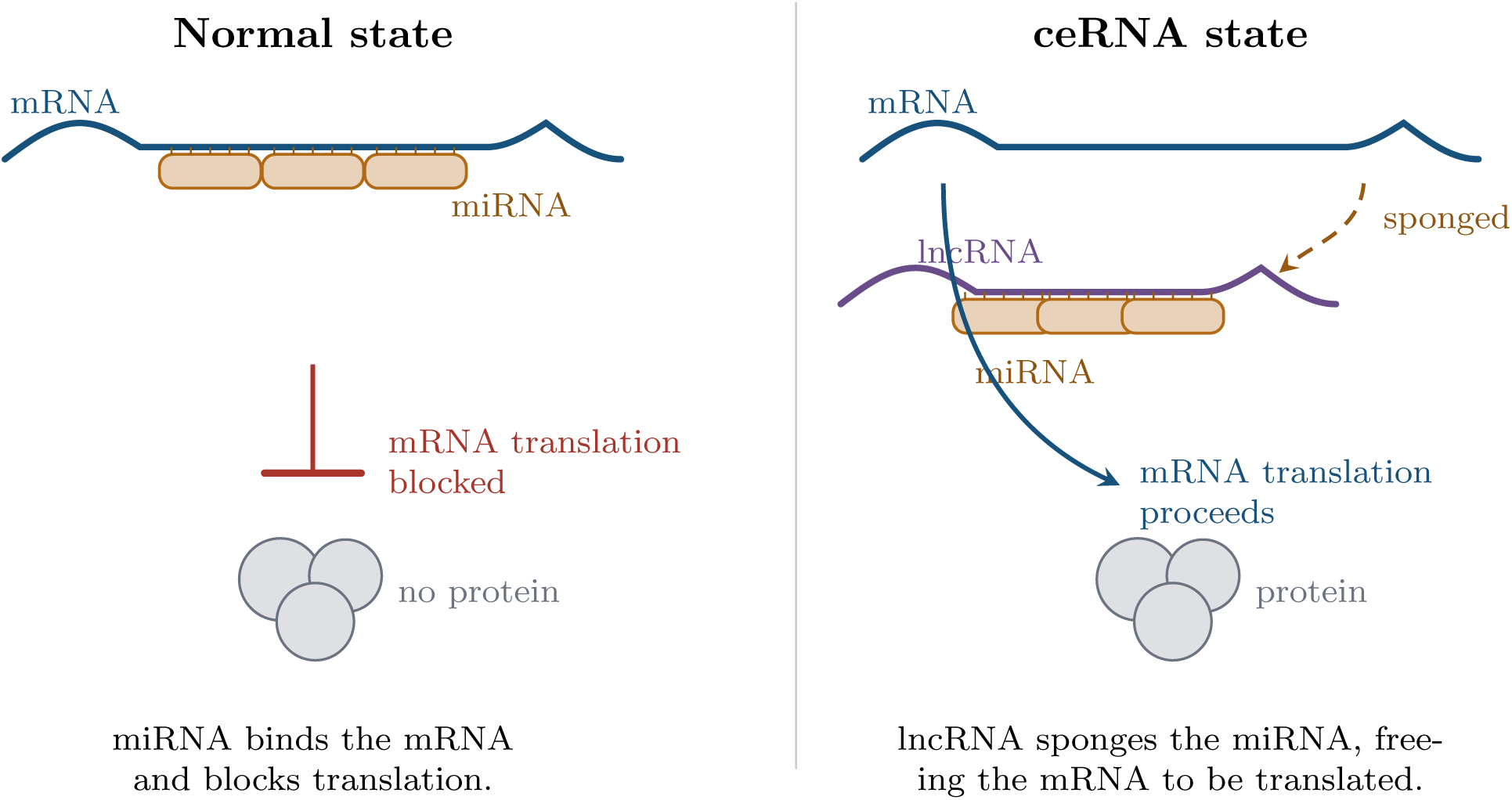
ceRNA theory. Left: miRNA binds the target mRNA and blocks its translation. Right: a long non-coding RNA competes for the same miRNA and sequesters it, so the mRNA, rather than the lncRNA, is released for translation. Colors follow Figure 2: mRNA blue, miRNA amber, lncRNA purple.

Recent research has shown that ceRNA interactions play an important role in glioblastoma. Su et al. (2021) confirmed the involvement of the lncRNA *MALAT1* in glioblastoma development: when *MALAT1* prevented the expression of miR-613, a miRNA implicated in glioblastoma, glioma cells proliferated more rapidly. Because such dysregulated RNAs contribute to glioblastoma progression, they are candidate molecular biomarkers, and a comprehensive analysis of ceRNA interactions offers an opportunity to identify new ones.

### 1.2 Graph Theory

A ceRNA network maps onto a heterogeneous graph, which admits more than one node and edge type. The network supplies three node (gene) types, miRNA, mRNA and lncRNA, and three edge (interaction) types: mRNA silencing, miRNA sponging and ceRNA co-regulation. Nodes additionally take features from the clinical metadata, stratified as detailed in Appendix A, so the broader medical context enters the representation. Treating RNA molecules as nodes and their interactions as distinct edge types therefore gives a tractable model of the network (Figure 2).

**Figure 2.**
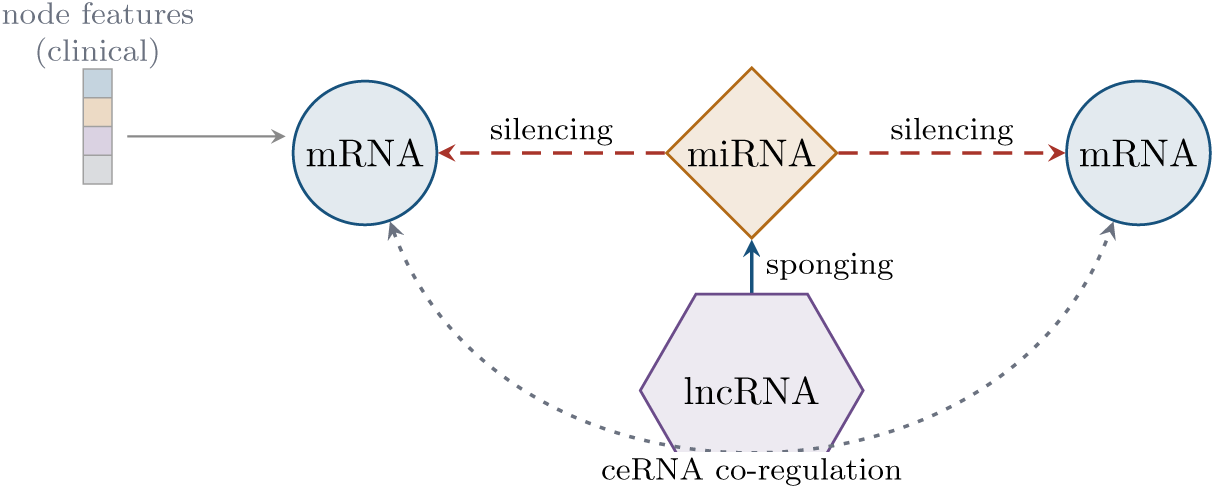
Heterogeneous representation of a ceRNA network. Shape and color both encode gene type: mRNA (circle), miRNA (diamond), lncRNA (hexagon). Edges show the three modeled interaction types: miRNA*→*mRNA silencing (dashed red), lncRNA*→*miRNA sponging (solid blue) and inferred ceRNA co-regulation between mRNAs sharing a miRNA regulator (dotted gray). Every node carries a vector of clinical features.

### 1.3 Machine Learning for ceRNA Network Analysis

RGCNs extend graph convolution to heterogeneous graphs by learning a separate transformation for each relation type (Schlichtkrull et al., 2017). Message passing builds each node’s representation from its neighbors and updates it recursively (Figure 3), so every gene is scored in the context of its surrounding interactions, which suits the structure of ceRNA data. Zhi et al. (2025) used the architecture to identify gastric cancer biomarkers, where it outperformed several established machine learning (ML) methods; that study attributed the margin to the model’s handling of distinct relationship types and of non-linear patterns.

**Figure 3.**
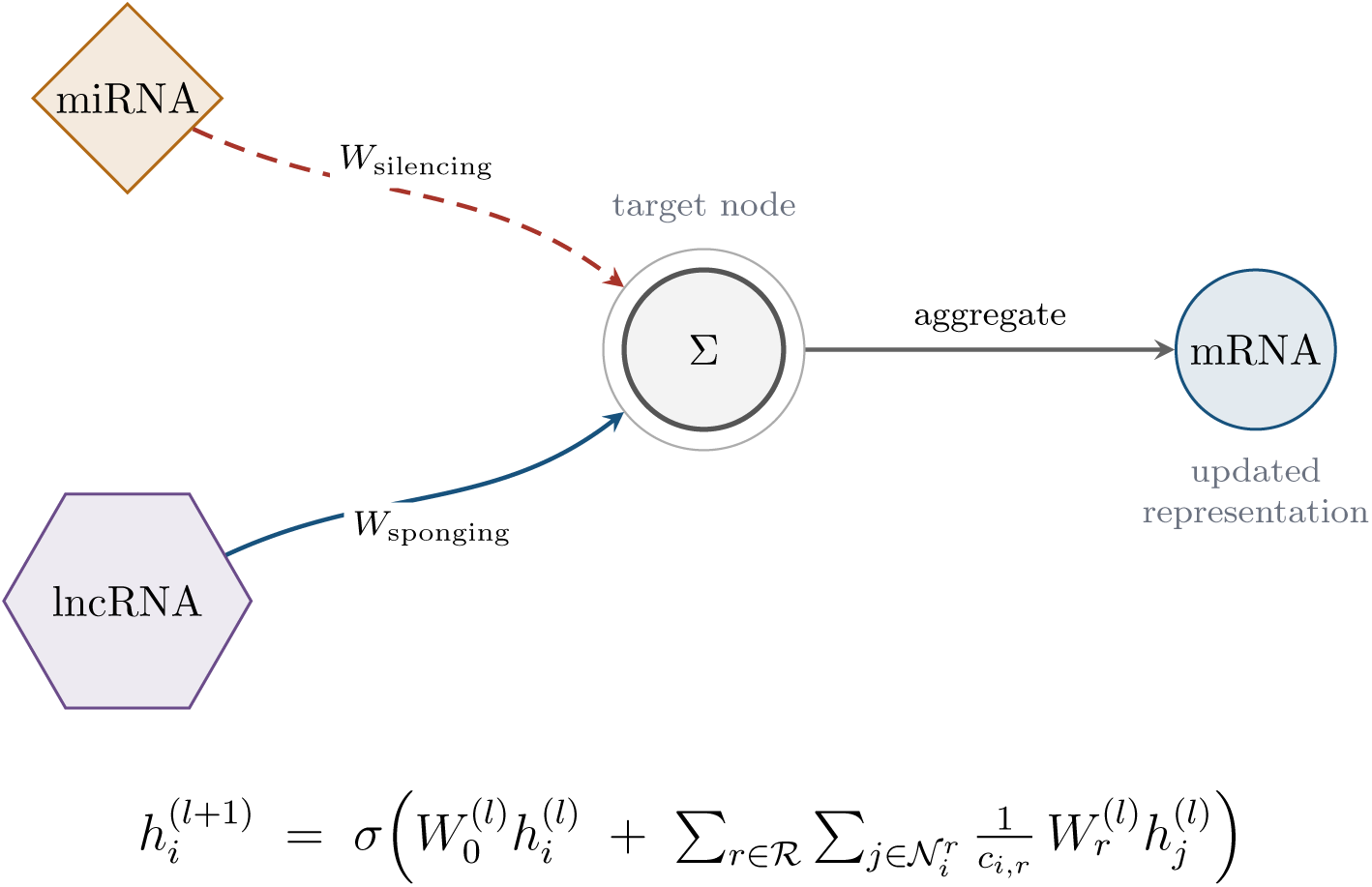
RGCN message passing. Each incoming relation is transformed by its own weight matrix (*W*_silencing_, *W*_sponging_) before the results are aggregated at the target node (Σ). In the layer update beneath, *h*^(^*^l^*^)^ is the representation of node *i* at layer *l*, *R* the set of relation types, *N ^r^* the neighbors of *i* under re*^i^*lation *r*, *c* a normalization constant, *W* the self-loop transformation, and *σ ^i^*the activation (Schlichtkrull et al., 2017).

### 1.4 Limitations of Current Statistical Approaches

Existing studies have applied various traditional ML and statistical methods to identify biomarkers via ceRNA network analysis. Liu et al. (2021) used weighted gene co-expression network analysis (WGCNA), which investigates the role of mRNA clusters by analyzing highly correlated genes. That study performed differential expression analysis, then functional enrichment analysis to narrow the candidates, and finally multivariate Cox regressions to evaluate each gene’s effect on survival, concluding that WGCNA could identify several genes associated with glioblastoma. However, WGCNA assumes simple, correlative relationships, which may miss biomarkers that participate in the extensive miRNA-mediated ceRNA interactions documented in glioblastoma (Sumazin et al., 2011).

Other approaches face similar limitations. Bazrgar et al. (2024) used a seed graph to construct a ceRNA network, applying differential expression analysis, using the most dysregulated genes as seed nodes, expanding the network using protein-protein interaction databases, and assessing survival associations to identify three prognostic regulatory axes. Although this approach benefits from building on genes already known to be significant, it relies heavily on the seed nodes and assumes that mRNA expression reflects protein expression. In glioblastoma, however, the correlation between an mRNA and its protein product can be low due to post-transcriptional processing such as miRNA silencing (Sumazin et al., 2011), producing statistical correlations between non-interacting genes. Deep learning architectures such as RGCNs can in principle weight relation types differently, but they do not by themselves distinguish direct interactions from indirect correlations, and we make no causal claim about the interactions modeled here.

### 1.5 Research Gap and Research Goals

The study by Zhi et al. (2025), which evaluated RGCNs for ceRNA analysis in gastric cancer, frames the research gap: RGCNs outperformed traditional ML and other graph neural network models in biomarker identification. Despite this, current analyses of glioblastoma ceRNA largely depend on statistical measures such as the WGCNA method of Liu et al. (2021). Correlation-based models are poorly suited to this setting, because glioblastoma is characterized by extreme lineage plasticity, with cancer cells transitioning between states depending on their environment, (Neftel et al., 2019), so the regulatory interactions active in a tumor are unlikely to be static. Because these models rely on static, linear associations, they can average out transient gene interactions and miss biomarkers that act through them. This paper addresses this gap by employing RGCNs to analyze the ceRNA network of glioblastoma.

## 2 MATERIALS AND METHODS

This study evaluated whether RGCNs can rank biomarker candidates in glioblastoma ceRNA networks, and whether integrating copy number variation (CNV) improves that ranking. The task is node-level classification: every node is scored on its likelihood of being a biomarker, and high-scoring nodes absent from the label set are treated as novel candidates. Models were compared by AUCROC, PR-AUC, and *F*_1_. The RGCN was tested against the thirteen alternatives in Table 1, spanning graph neural baselines, non-graph baselines, and the CNV fusion arms, alongside three reference baselines that fix the floor for each metric. All analyses were performed in Python and are publicly available. An overview of the analysis pipeline is shown in Figure 4.

**Figure 4.**
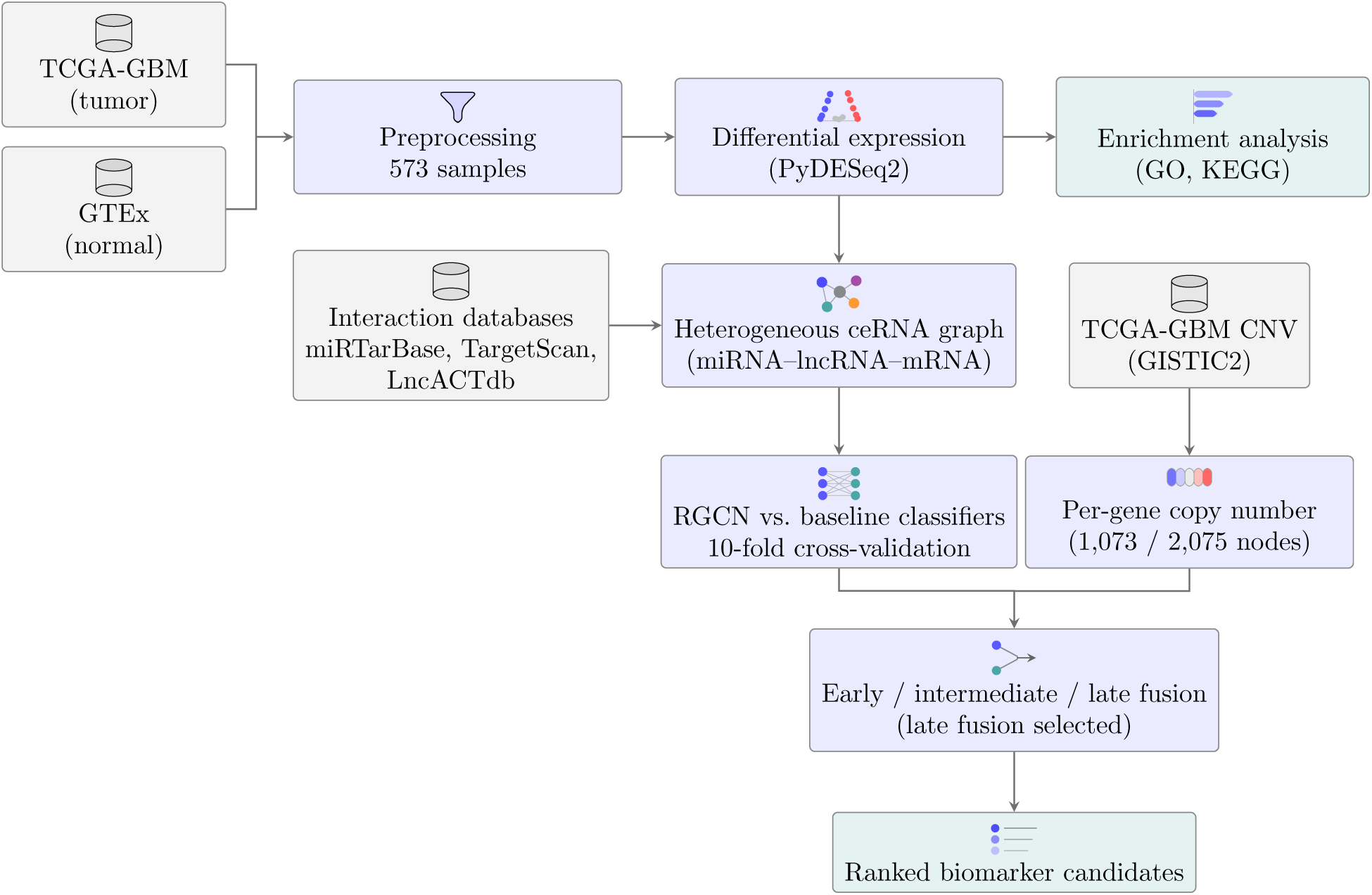
Overview of the analysis pipeline. Differential expression on the tumor and normal data yields both the enrichment gene set and, with curated interaction databases, the heterogeneous ceRNA graph; GISTIC2 calls supply a per-gene copy-number score in parallel. Models are scored under 10-fold cross-validation, and the two branches are combined by early, intermediate or late fusion, of which late fusion performed best.

**Table 1.** Model performance across all evaluated architectures, reported as mean *±* SD over 10-fold cross-validation. Models are grouped into graph neural networks trained on the ceRNA graph, non-graph baselines, and the CNV fusion architectures. Bold marks the best value for each metric within its group.

| Model | AUCROC | PR-AUC | F1 | Accuracy | Precision | Recall |
| --- | --- | --- | --- | --- | --- | --- |
| *Graph neural networks (ceRNA graph)* |  |  |  |  |  |  |
| RGCN | **0.874 ± 0.070** | **0.124 ± 0.061** | 0.054 ± 0.080 | **0.904 ± 0.094** | 0.032 ± 0.050 | 0.283 ± 0.416 |
| RGAT | 0.794 ± 0.110 | 0.078 ± 0.057 | **0.109 ± 0.080** | 0.742 ± 0.260 | **0.062 ± 0.047** | **0.675 ± 0.352** |
| GAT | 0.673 ± 0.186 | 0.074 ± 0.100 | 0.046 ± 0.039 | 0.742 ± 0.257 | 0.025 ± 0.022 | 0.417 ± 0.345 |
| GCN | 0.545 ± 0.205 | 0.071 ± 0.102 | 0.042 ± 0.040 | 0.735 ± 0.275 | 0.024 ± 0.024 | 0.325 ± 0.315 |
| *Non-graph baselines* |  |  |  |  |  |  |
| Hierarchical NN | **0.856 ± 0.111** | 0.145 ± 0.082 | 0.138 ± 0.051 | 0.876 ± 0.057 | 0.080 ± 0.033 | 0.617 ± 0.249 |
| Random forest | 0.818 ± 0.138 | 0.142 ± 0.068 | **0.173 ± 0.171** | **0.960 ± 0.021** | **0.150 ± 0.173** | 0.267 ± 0.263 |
| k-NN | 0.772 ± 0.140 | **0.151 ± 0.129** | 0.118 ± 0.101 | 0.919 ± 0.055 | 0.084 ± 0.097 | 0.358 ± 0.294 |
| Node2Vec + LR | 0.725 ± 0.119 | 0.055 ± 0.028 | 0.036 ± 0.027 | 0.666 ± 0.249 | 0.019 ± 0.014 | 0.567 ± 0.446 |
| Logistic regression | 0.738 ± 0.073 | 0.042 ± 0.015 | 0.063 ± 0.033 | 0.699 ± 0.130 | 0.034 ± 0.018 | **0.675 ± 0.315** |
| *CNV fusion architectures* |  |  |  |  |  |  |
| RGCN (ceRNA only)<sup>†</sup> | 0.867 ± 0.064 | 0.109 ± 0.046 | 0.081 ± 0.071 | 0.873 ± 0.097 | 0.047 ± 0.044 | 0.450 ± 0.416 |
| CNV only | 0.698 ± 0.115 | 0.027 ± 0.007 | 0.052 ± 0.013 | 0.521 ± 0.038 | 0.027 ± 0.007 | **0.875 ± 0.227** |
| Early fusion | 0.853 ± 0.092 | 0.111 ± 0.044 | 0.076 ± 0.106 | **0.928 ± 0.069** | 0.048 ± 0.070 | 0.267 ± 0.378 |
| Intermediate fusion | 0.860 ± 0.098 | 0.137 ± 0.114 | 0.073 ± 0.090 | 0.901 ± 0.087 | 0.044 ± 0.056 | 0.325 ± 0.417 |
| Late fusion | **0.883 ± 0.072** | **0.208 ± 0.152** | **0.132 ± 0.109** | 0.911 ± 0.073 | **0.091 ± 0.098** | 0.442 ± 0.360 |
| *Reference baselines* |  |  |  |  |  |  |
| All negative | 0.500 | 0.015 | 0.000 | 0.985 | 0.000 | 0.000 |
| Random ranker | 0.500 | 0.015 | — | — | 0.015 | — |
| Node type only<sup>‡</sup> | 0.824 ± 0.139 | 0.075 ± 0.025 | 0.150 ± 0.035 | 0.859 ± 0.023 | 0.083 ± 0.020 | 0.842 ± 0.231 |

<sup>†</sup>The two RGCN rows are the same architecture evaluated under two protocols: the row in the first group is trained on the ceRNA graph with the original node features, while the fusion-arm row is the ceRNA-only control for the fusion experiment and uses the fusion feature matrix and fold assignment. They are not independent models and should not be read as two results. <sup>‡</sup>Logistic regression on one-hot node type with no other input (Section 3.2). Reference rows are not bolded; the random ranker's threshold-dependent metrics depend on the cut-off and are omitted.

The proposed framework consists of two primary branches. The ceRNA branch encodes post-transcriptional regulation through a heterogeneous graph, consisting of miRNA, lncRNA, and mRNA nodes joined by three edge types. The CNV branch represents genomic dosage as a per-gene copy-number score. Three forms of fusion were tested: early, intermediate, and late, combining the two branches at the feature, representation and decision level respectively.

### 2.1 Data Preparation and Processing

Diseased-tissue gene expression data were obtained from the TCGA-GBM (The Cancer Genome Atlas Glioblastoma) dataset via the UCSC Xena Toil Recompute Project (The Cancer Genome Atlas Research Network, 2008; Goldman et al., 2020). TCGA-GBM contains RNA sequencing data from tumor samples for mRNA, lncRNA, and miRNA. It was selected over alternative datasets (e.g., GSE90604) because of its large sample size and comprehensive clinical metadata; however, it consists largely of patients from Western populations, which limits its generalizability.

TCGA-GBM provides little normal tissue, which differential expression analysis requires, so normal control data were taken from the Genotype-Tissue Expression (GTEx) dataset (The GTEx Consortium, 2013). GTEx carries no miRNA profiles, so the few normal samples in TCGA-GBM had to serve as the miRNA controls, which likely weakened the statistical strength of some model metrics. Both resources were chosen for their standardized protocols, coverage, and public availability, and for the clinical metadata that supplies the node features described below. Liu et al. (2021) and Bazrgar et al. (2024) draw on the same sources.

#### 2.1.1 Preprocessing

Of the 612 available samples, 14 duplicates, 11 low-quality, and 14 outliers were removed, leaving 573 samples for analysis. Outliers were identified with PyDESeq2 (Muzellec et al., 2023); samples were counted as low-quality if they lacked clinical metadata. All missing gene expression values were set to 0, and negative values were clipped to 0. The Toil recompute distributes RSEM expected counts, which are non-integer because reads are apportioned probabilistically across isoforms; these were rounded to the nearest integer for compatibility with the negative-binomial model used by DESeq2. miRNA expression was analyzed from raw read counts, as described below. Gene type classifications were obtained from the MyGene.info API v3 (Wu et al., 2013).

miRNA read counts were taken from the TCGA-GBM miRNA-seq quantification files of the NCI Genomic Data Commons (`read_count` column), keeping one aliquot per sample. Only five solid-tissue normal samples are available, all sequenced in the same batch as six primary tumors; the miRNA contrast therefore compares these 6 tumor and 5 normal samples, not the full 573-sample cohort, which applies only to the mRNA and lncRNA arms. The graph was built from an earlier version of this contrast computed on the Xena GDC matrix, which is distributed on a log_2_(*x* + 1) scale and had been converted back to pseudo-counts; re-estimation on raw counts agreed closely with it (Pearson *r* = 0.989 between log_2_ fold changes) and retained 285 of the 314 graph miRNAs, so the graph was not rebuilt. With DESeq2 fold-change shrinkage, the candidate effect sizes in Section 3.3 were essentially unchanged. As a sensitivity analysis, contrasting all 267 available primary tumors against the same five normals reproduced the direction and significance of all five candidate loci, although that comparison confounds tumor status with sequencing batch. With five controls the variance of the normal baseline remains poorly estimated, and we regard the miRNA arm as exploratory on that basis.

#### 2.1.2 Differential Expression Analysis

Differential expression was computed with PyDESeq2 0.5.4 (Muzellec et al., 2023), controlling the false discovery rate at a Benjamini–Hochberg adjusted *p*-value *<* 0.05 with *|* log_2_ fold change*| >* 1. This yielded 1,166 differentially expressed mRNA, 595 differentially expressed lncRNA, and 314 differentially expressed miRNA, giving 2,075 genes in total. Functional enrichment analysis was then run on that set as a check on its coherence (Appendix E); the recovered pathways align with those reported in existing studies, which supports the biological relevance of the processed dataset (Bazrgar et al., 2024; Liu et al., 2021).

### 2.2 Heterogeneous Graph Construction

Constructing a graph requires three key components: nodes (genes), node features (clinical data), and edges (interactions). The nodes were obtained from the differential expression analysis above. To generate the node features, an independent DE analysis was performed on the clinical features across all genes, using the stratification criteria listed in Appendix A. The resulting per-gene log_2_ value became a node feature, allowing the model to capture how a gene is differentially expressed across each clinical axis. Incorporating these clinical features enables the model to determine which genes are associated with specific clinical profiles and how those profiles influence their interactions with other ceRNAs, which matters because glioblastoma is molecularly heterogeneous across patient subgroups (Verhaak et al., 2010). Zhi et al. (2025) construct node features in a similar way. Each node therefore carries a ten-dimensional feature vector: nine clinical axes (Appendix A) and the copy-number channel described below.

The resulting graph contains 2,075 nodes (1,166 mRNA, 595 lncRNA, and 314 miRNA) joined by 4,070 directed edges across three relation types. Of these, 1344 are curated from interaction databases: 1030 miRNA*→*mRNA silencing edges (25.3% of all edges) and 314 lncRNA*→*miRNA sponging edges (7.7%), drawn from miRTarBase (Huang et al., 2022) and LncACTdb (Wang et al., 2022), supplemented by TargetScan (Agarwal et al., 2015) predictions retained above a context score of 0.6. ENCORI was queried but returned no records passing filtering, and contributes no edges.

The remaining 2,726 edges (67.0%) are ceRNA co-regulation edges, which are inferred rather than curated. An undirected pair of edges is added between two mRNAs whenever they share at least one common miRNA regulator among the curated silencing edges. The rule is deliberately permissive: it sets no threshold on the number of shared regulators and applies no expression-correlation filter, so hub miRNAs with many targets generate dense cliques among their targets. Because this inference supplies the majority of the graph, model performance is conditional on it; a stricter rule requiring several shared regulators, or agreement in expression, would give a sparser graph and potentially different rankings. We report this rule explicitly because the controls in Section 3.2 show that the relation structure encodes node type, and this rule is what makes mRNA–mRNA adjacency dense relative to the other two relations.

### 2.3 Copy Number Variation Data

Copy-number data were obtained from the TCGA-GBM GISTIC2 (Mermel et al., 2011) thresholded gene-level calls distributed through UCSC Xena, covering 24,776 genes across 577 samples. GISTIC2 reports integer calls in *{−*2*, −*1, 0, +1, +2*}*, corresponding to deep deletion, shallow deletion, copy-neutral, gain, and amplification. Per-gene scores were averaged across samples and mapped onto graph nodes by HGNC symbol, resolving 86.4% of symbols.

Coverage is uneven across node types, and the fusion results below turn on that unevenness. Copy-number values were resolved for 1073 of the 2,075 nodes (51.7%): 1013 of 1,166 mRNA nodes (86.9%) and 60 of 595 lncRNA nodes (10.1%), but none of the 314 miRNA nodes, since GISTIC2 segment boundaries are undefined at miRNA loci. Each node therefore carries a binary `cnv observed` flag alongside its copy-number value, keeping an unmeasured node distinguishable from a genuinely copy-neutral one. The mapping checks out against known biology: the most amplified gene genome-wide is *EGFR* (+1.307) and the most deleted is *CDKN2A* (*−*1.296), both canonical glioblastoma events.

### 2.4 Model Architecture and Training

The RGCN comprises two `RGCNConv` layers, each maintaining a separate transformation per relation type, with a hidden and output dimension of 64, ReLU activation, and dropout at *p* = 0.3 between layers. Node representations are passed to a two-layer classification head (64 *→* 64 *→* 1) producing a single logit per node. The graph baselines (GCN, GAT, RGAT) use the same hidden dimensions, dropout, and head, differing only in their convolution operator; GAT and RGAT use four attention heads in the first layer.

Models were trained with Adam (learning rate 10*^−^*^3^, weight decay 10*^−^*^4^) for up to 100 epochs, minimizing binary cross-entropy with logits. Because only 31 of the 2,075 nodes carry a positive label, the positive class was weighted by the negative-to-positive ratio, giving a `pos_weight` of approximately 65.9. Training used early stopping on validation AUCROC with a patience of 10 epochs, and the parameters from the best validation epoch were restored before testing.

### 2.5 Evaluation Protocol

All models were evaluated by stratified 10-fold cross-validation under a fixed random seed, so every architecture was scored on identical folds. Within each training fold a further stratified 10% split was held out for validation, used both for early stopping and for selecting the decision threshold; test folds were never consulted during training or model selection. Because the class balance makes a fixed 0.5 cut-off uninformative, the threshold was chosen to maximize validation *F*_1_ over a grid from 0.05 to 0.95, and then applied unchanged to the test fold. Threshold-free metrics (AUCROC, PR-AUC) are unaffected by this choice; the threshold-dependent metrics (accuracy, precision, recall, and *F*_1_) reflect a tuned operating point.

Given the 1.49% positive rate, PR-AUC is the more informative summary: a random ranker achieves 0.015, whereas the AUCROC of such a ranker is 0.5 regardless of prevalence (Saito and Rehmsmeier, 2015). At this prevalence accuracy is dominated by the negative class — a model that labels every node negative scores 0.985 — and precision must be read against its chance level of 0.015, so Table 1 includes all-negative, random, and node-type-only reference rows. For the controls we additionally report the Matthews correlation coefficient (MCC) and balanced accuracy, which, unlike accuracy and *F*_1_, are high only when performance is good on both classes (Chicco and Jurman, 2020). Model pairs were compared by the paired Wilcoxon signed-rank test across the ten folds, which respects the pairing induced by the shared fold assignment.

The evaluation is transductive: the graph is fixed, message passing runs over all 2,075 nodes at every epoch, and the folds partition the labels rather than the graph, so a test node’s neighborhood is visible during training even though its label is withheld. Reported figures are therefore transductive node-classification estimates and not an estimate of performance on nodes added to the graph later. Because ten folds of three to four positives each give very coarse per-fold metrics, we additionally report percentile bootstrap confidence intervals on the pooled out-of-fold predictions, which use all 31 positives at once.

### 2.6 CNV Fusion Strategies

Three fusion strategies were compared against the ceRNA-only RGCN. *Early fusion* appends the copy-number value and its observation flag to the node feature vector, leaving a single graph stream. *Intermediate fusion* adds a parallel multilayer perceptron over the copy-number channel whose output is concatenated with the graph embedding before the classification head. *Late fusion* trains the two branches independently, the RGCN on the ceRNA graph and a balanced random forest on the tabular copy-number features, then blends their predicted probabilities as *αp*_graph_ + (1 *− α*)*p*_CNV_. The mixing weight *α* was selected per fold on the validation split by maximizing validation AUCROC over a grid from 0 to 1, and never on test data.

## 3 RESULTS

### 3.1 Model Performance

The RGCN and all baseline models were trained and tested on the same processed dataset. Among the models trained on the ceRNA graph alone, the RGCN discriminated best, reaching a mean AUCROC of 0.874 *±* 0.070 across 10-fold cross-validation with an accuracy of 0.904 *±* 0.094. Under a paired Wilcoxon signed-rank test across folds it outperformed GCN (Δ = +0.328, *p* = 0.004), GAT (Δ = +0.201, *p* = 0.027), RGAT (Δ = +0.079, *p* = 0.027) and logistic regression (Δ = +0.136, *p* = 0.006); Table 2 gives the full comparison. Its low precision (0.032 *±* 0.050) indicated a high false-positive rate, a critical bottleneck requiring downstream *in vitro* filtering, although its PR-AUC of 0.124 is roughly eight times the chance level of 0.015. The corresponding ROC curves are shown in Figure 5.

**Figure 5.**
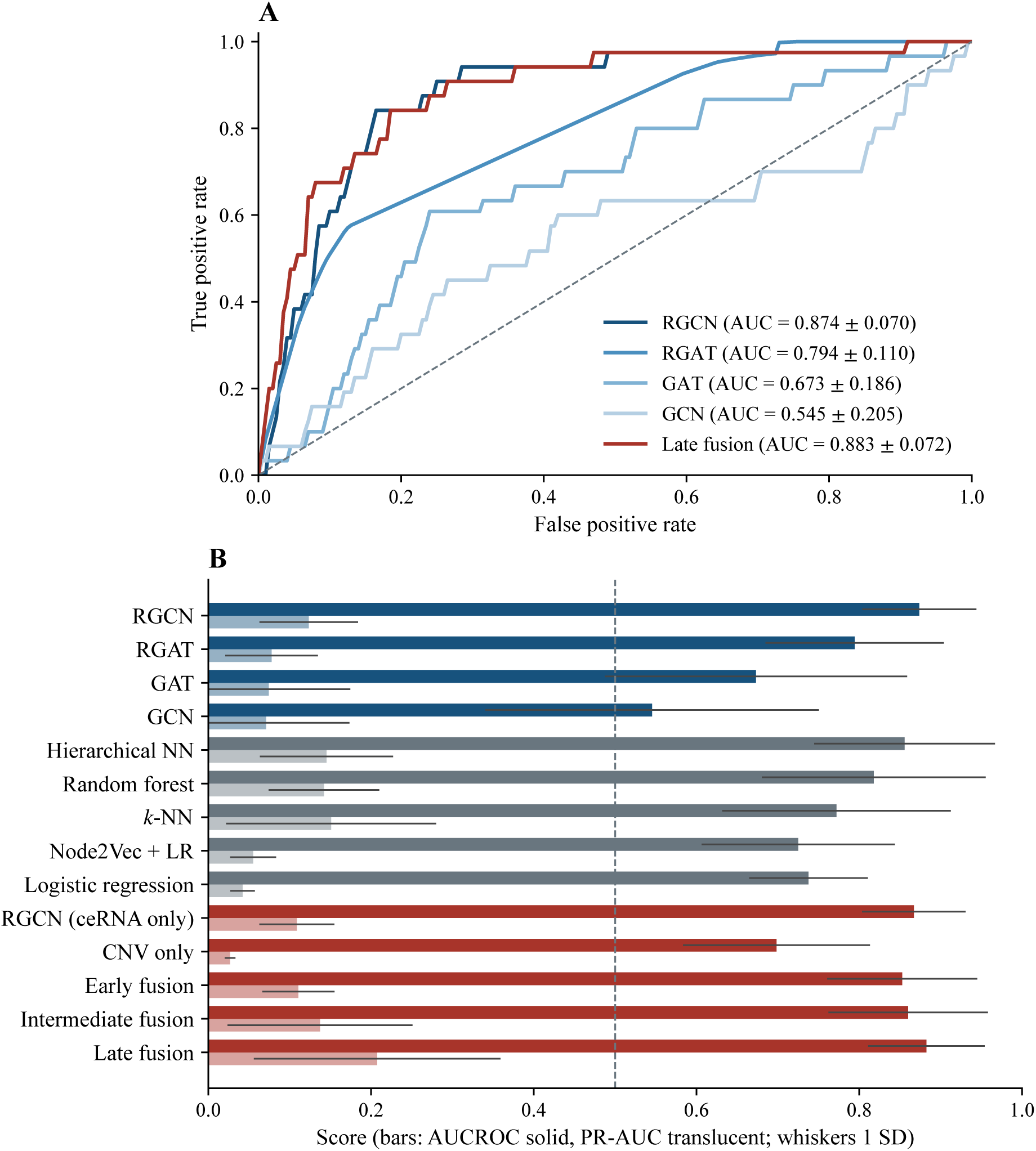
ROC curve comparison for GBM biomarker classification. The diagonal is the random-classifier baseline. Curves are the mean true-positive rate across the ten cross-validation folds, and the reported AUC values are the corresponding fold means *±* standard deviation.

**Table 2.** Pairwise significance testing between selected model pairs, using the paired Wilcoxon signed-rank test across the ten cross-validation folds. Δ is the mean difference in the stated metric, with positive values favoring the first-named model.

| Comparison | Metric | $\Delta$ | $W$ | $p$ |
| --- | --- | --- | --- | --- |
| RGCN vs. GCN | AUCROC | +0.3283 | 1.0 | 0.0039* |
| RGCN vs. GAT | AUCROC | +0.2006 | 6.0 | 0.0273* |
| RGCN vs. RGAT | AUCROC | +0.0794 | 6.0 | 0.0273* |
| RGCN vs. Logistic regression | AUCROC | +0.1361 | 2.0 | 0.0059* |
| Late fusion vs. RGCN (ceRNA only) | AUCROC | +0.0155 | 7.0 | 0.1484 |
| Late fusion vs. RGCN (ceRNA only) | PR-AUC | +0.0987 | 0.0 | 0.0078* |
| Late fusion vs. Intermediate fusion | PR-AUC | +0.0702 | 4.0 | 0.0137* |
| Late fusion vs. Early fusion | PR-AUC | +0.0968 | 7.0 | 0.0371* |
| Late fusion vs. CNV only | PR-AUC | +0.1807 | 0.0 | 0.0020* |
| Intermediate fusion vs. RGCN (ceRNA only) | PR-AUC | +0.0285 | 22.0 | 0.6250 |
| Early fusion vs. RGCN (ceRNA only) | PR-AUC | +0.0019 | 27.0 | 1.0000 |

\*Significant at $\alpha = 0.05$ . Reported $p$ -values are uncorrected. Under Holm–Bonferroni correction across all eleven comparisons, two remain significant: late fusion versus CNV-only on PR-AUC ( $p_{\text{Holm}} = 0.022$ ) and RGCN versus GCN on AUCROC ( $p_{\text{Holm}} = 0.039$ ). The remainder, including late fusion versus the ceRNA-only RGCN ( $p_{\text{Holm}} = 0.063$ ), do not reach significance under this stricter criterion.

Integrating the CNV branch improved performance further. The late-fusion ensemble was the strongest model overall, with a mean AUCROC of 0.883 *±* 0.072 and a mean PR-AUC of 0.208 *±* 0.152. Only 31 of the 2,075 scored nodes carry a positive label, so PR-AUC is the more informative metric here (a random ranker attains 0.015), and the benefit of fusion is clearest on it. Before correction for multiple comparisons, late fusion outperformed the ceRNA-only RGCN (Δ = +0.099, *p* = 0.008), the CNV-only model (Δ = +0.181, *p* = 0.002), and both alternative fusion strategies: intermediate (Δ = +0.070, *p* = 0.014) and early (Δ = +0.097, *p* = 0.037). Neither early nor intermediate fusion improved significantly on the ceRNA-only baseline (*p* = 1.000 and *p* = 0.625 respectively), which points to the decision-level combination itself as the source of the advantage, not the addition of copy-number information. These *p*-values are uncorrected; under Holm–Bonferroni correction across the eleven comparisons in Table 2, only the late fusion versus CNV-only and RGCN versus GCN comparisons remain significant, and the late fusion advantage over the ceRNA-only RGCN falls just outside the threshold (*p*_Holm_ = 0.063). Ten folds give the test limited power, so the evidence for decision-level fusion is suggestive and not conclusive.

### 3.2 Controls: Node-Type Prior and CNV Observability

The positive labels are not distributed evenly across node types. Of the 31 labeled positives, 26 are miRNA, so 8.28% of miRNA nodes carry a positive label against 0.34% of lncRNA and 0.26% of mRNA nodes — a miRNA node is roughly 32 times more likely to be positive than an mRNA node. Node type is recoverable from the relation structure, because the three relation types connect the three node types in a fixed pattern. A model could therefore score well by learning node type rather than anything about ceRNA topology. Three controls separate the two explanations; all use the evaluation protocol of Section 2.5 and are reported in Table 3. The controls were run together with a fresh fit of the reference RGCN in a single software environment, in which it reached an AUCROC of 0.876 *±* 0.069 and a PR-AUC of 0.154 *±* 0.105; comparisons below use that run.

**Table 3.** Controls separating ceRNA topology from the node-type prior and from CNV observability. All values use the 10-fold protocol of Section 2.5; CIs are percentile bootstrap intervals on pooled out-of-fold predictions.

| Control | AUCROC (mean ± SD) | PR-AUC | Pooled AUCROC [95% CI] |
| --- | --- | --- | --- |
| *Node-type prior* |  |  |  |
| RGCN, full graph (reference) | 0.876 ± 0.069 | 0.154 | — |
| Node type only, no topology | 0.824 ± 0.139 | 0.075 | 0.790 [0.670, 0.890] |
| RGCN, scored on miRNA nodes | 0.613 ± 0.173 | 0.196<sup>a</sup> | 0.568 [0.468, 0.663] |
| RGCN, trained on miRNA nodes only | 0.432 ± 0.180 | — | 0.469 [0.345, 0.593] |
| Within-type permutation null<sup>b</sup> | 0.799 ± 0.042 | 0.142 ± 0.042 | — |
| *CNV observability* |  |  |  |
| (1 - cnv_observed) alone | 0.697 | — | — |
| CNV value only, observed nodes | 0.475 ± 0.007 | — | 0.472 [0.464, 0.479] |

<sup>a</sup>Chance level within miRNA nodes is 0.083. <sup>b</sup>Mean ± SD over 200 permutations, with the RGCN retrained on each; empirical $p = 0.015$ (AUCROC) and $p = 0.33$ (PR-AUC) for the observed reference. The miRNA-only and CNV-value-only arms carry 26 and 4 positives respectively; the latter is underpowered and is reported for completeness rather than as evidence of absence.

*A node-type-only baseline*, a logistic regression on a one-hot node-type vector and no other information, reaches a mean AUCROC of 0.824 *±* 0.139 (pooled out-of-fold 0.790, 95% bootstrap CI [0.670, 0.890]), within the fold-to-fold variation of the RGCN. On PR-AUC the type-only model reaches 0.075 *±* 0.025, about half the RGCN’s value, while on the threshold-dependent summaries it scores higher than the RGCN (MCC 0.208 against 0.133; balanced accuracy 0.823 against 0.678). Most of the reported discrimination is therefore obtainable with no topological information.

*Restricting evaluation to the 314 miRNA nodes*, where the type prior is constant by construction, gives weak discrimination. Scoring the full-graph RGCN on its miRNA test nodes gives an AUCROC of 0.613 *±* 0.173 and a PR-AUC of 0.196 against a chance level of 0.083, but the pooled estimate, 0.568 (CI [0.468, 0.663]), does not exclude chance; an RGCN trained and evaluated on the miRNA subgraph alone reaches 0.432 *±* 0.180, with seven of ten folds below 0.5.

*Permuting labels within node type* preserves the prior exactly while destroying any real topological signal. We retrained the RGCN under the full cross-validation protocol on each of 200 such permutations (Figure 6). On PR-AUC the observed value is indistinguishable from this null (0.154 against 0.142 *±* 0.042, empirical *p* = 0.33), as it is on MCC (*p* = 0.35): the RGCN’s PR-AUC advantage over the logistic type-only model reflects more efficient use of the type prior, not topology. On AUCROC the observed value exceeds the null (0.876 against 0.799 *±* 0.042, *p* = 0.015), indicating a modest signal beyond node type; within the miRNA subgraph that signal does not reach significance (*p* = 0.10).

**Figure 6.**
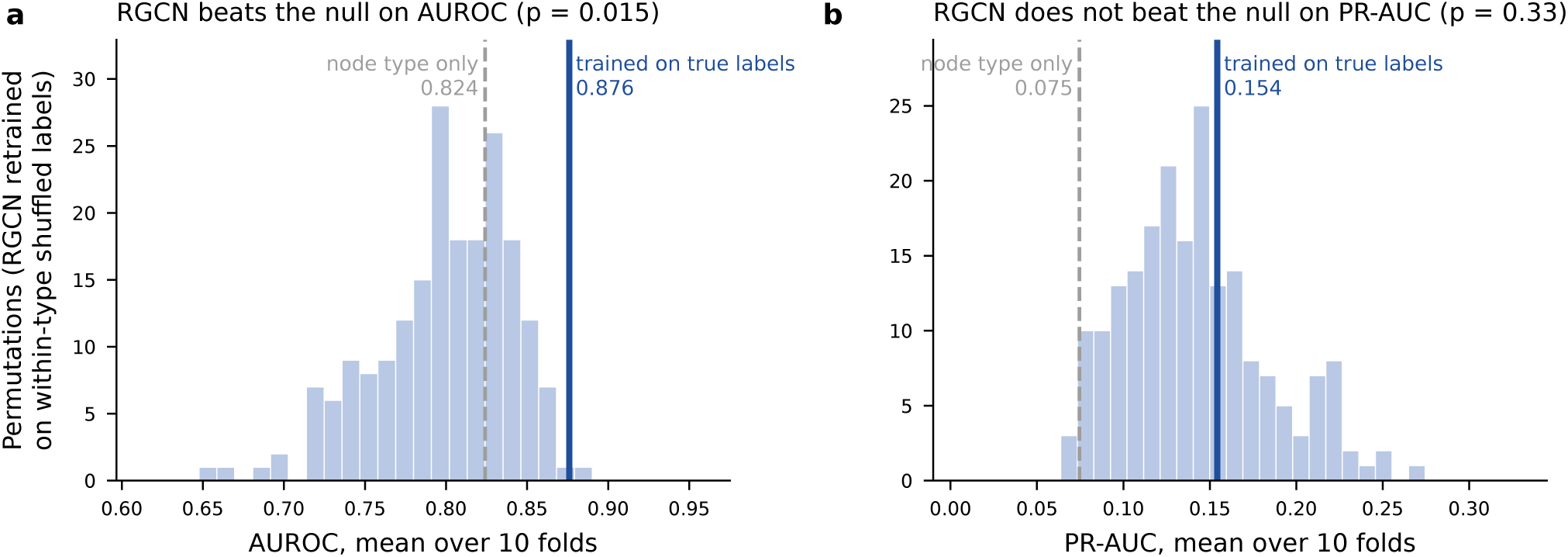
Within-type permutation null with the RGCN retrained on each of 200 permutations. Histograms show the null distribution of the 10-fold mean AUCROC (a) and PR-AUC (b); the solid line marks the RGCN trained on the true labels and the dashed line the node-type-only logistic regression.

The fusion result has the same origin. Copy number is undefined at all 314 miRNA loci, so the binary `cnv_observed` flag is close to a miRNA indicator: 27 of the 31 positives sit on nodes with no copy-number measurement. Used as a lone predictor, (1 *−* `cnv_observed`) reaches an AUCROC of 0.697, which reproduces the CNV-only row of Table 1 (0.698 *±* 0.115) almost exactly. Retraining the CNV branch on the 1,073 CNV-observed nodes with the flag dropped, so that it sees only the copy-number value, gives 0.475 *±* 0.007 — indistinguishable from chance. On that same subset the copy-number value adds +0.067 AUCROC over the graph branch even when the mixing weight is tuned on the test fold, an optimistic upper bound. We note that only 4 of the 31 positives survive the restriction to CNV-observed nodes, so this arm is severely underpowered and we do not read a negative result from it; what it does establish is that the CNV-only performance reported in Table 1 is carried by observability, not by genomic dosage.

*Imbalance-aware training* does not change this picture. Varying the positive-class weight (half and double the default), replacing the loss with a focal loss (*γ ∈ {*1, 2, 3*}*, *α ∈ {*0.5, 0.75, 0.9*}*), or treating unlabeled nodes as unlabeled rather than negative with a non-negative positive-unlabeled loss (class prior 0.015–0.10), each selected on validation PR-AUC, gave no improvement over the reference RGCN: the selected focal and positive-unlabeled variants reached PR-AUC values of 0.077 and 0.099, against 0.154. Choosing the decision threshold by validation MCC rather than *F*_1_ left the RGCN’s MCC essentially unchanged (0.133 against 0.129). Performance on this label set is limited by its composition rather than by the training objective (Appendix D).

### 3.3 Novel Biomarkers

Candidates were taken from the late-fusion ensemble, the best-performing configuration reported above. Because the aim is to nominate targets that prior work has not already characterized, the ranking was screened for existing glioma literature: for each locus we counted PubMed records naming the miRNA together with glioma, glioblastoma or GBM in the title or abstract, and retained those with at most one such record. The screen matches each locus under its miRBase precursor identifier and its conventional naming variants — the mature -3p and -5p strands, the bare numeric form, and, for numbered locus copies such as hsa-mir-450a-2, the copy-independent stem under which the shared mature product is named — and takes the union of matching records. Alias expansion matters: searching precursor identifiers alone returns 3, 1 and 0 records for the three well-characterized loci below, against 19, 12 and 11 once variants are included. Several of the highest-ranked loci are already well represented in the glioma literature (hsa-mir-196a, 19 records; hsa-mir-224, 12; hsa-mir-135b, 11) and were therefore not carried forward as novel candidates.

Applying this screen down the ranking gives five loci, none of them indexed in the curated glioblastoma biomarker set used for labeling and none the subject of more than one prior glioma report under alias-expanded search: hsa-mir-1258 (1 record), hsa-mir-203b (0), hsa-mir-450a-2 (1), hsa-mir-888 (1), and hsa-mir-5683 (0). Expansion changed two of these counts relative to a precursor-only search, which found no records for hsa-mir-1258 or hsa-mir-450a-2; both are matched only through their mature or copy-independent names. Three are downregulated in tumor tissue and two upregulated, with *|* log_2_ fold change*|* from 5.64 to 7.55 (all Benjamini–Hochberg adjusted *p <* 0.05). Table 4 lists them with their differential-expression statistics, including assay abundance and the standard error of each effect estimate, and their composite confidence scores, which range from 9.44 to 7.84 against 16.24 for the highest-ranked locus overall.

**Table 4.** Novel candidate biomarker loci from the late-fusion RGCN ensemble, after screening out loci with existing glioma literature. Rank is the position in the unscreened late-fusion ranking; prior reports is the count of PubMed records naming the miRNA together with glioma, glioblastoma or GBM in the title or abstract.

| Rank | miRNA locus | Base mean | log<sub>2</sub>FC | lfcSE | Adj. *p* | Confidence | Prior reports |
| --- | --- | --- | --- | --- | --- | --- | --- |
| 8 | hsa-mir-1258 | 56.8 | −5.645 | 1.099 | $1.41 \times 10^{-6}$ | 9.44 | 1 |
| 10 | hsa-mir-203b | 186.7 | −6.031 | 0.752 | $1.38 \times 10^{-14}$ | 8.58 | 0 |
| 12 | hsa-mir-450a-2 | 847.8 | +6.043 | 0.573 | $2.13 \times 10^{-24}$ | 8.48 | 1 |
| 13 | hsa-mir-888 | 65.3 | +7.546 | 1.359 | $1.61 \times 10^{-7}$ | 7.91 | 1 |
| 15 | hsa-mir-5683 | 179.0 | −6.945 | 0.846 | $3.22 \times 10^{-15}$ | 7.84 | 0 |

Base mean is the DESeq2 normalized mean count across all samples and indicates assay abundance; lfcSE is the standard error of the log<sub>2</sub> fold change. The two lowest-abundance loci are hsa-mir-1258 and hsa-mir-888 (base mean 56.8 and 65.3); hsa-mir-888 additionally carries the largest standard error (1.359) and is detected in only one normal sample, so its effect estimate is the least secure of the five. All contrasts are estimated from raw read counts on 6 tumor against 5 normal samples (Section 2.1.1). Confidence is a composite of the cross-validated node score, the absolute log<sub>2</sub> fold change, and eigenvector centrality, combined on a 0–20 scale; it is a ranking device internal to this study and is not calibrated or comparable across studies. Given Section 3.2, it should be read as descriptive rather than as evidence of biomarker status. Hazard ratios are not reported, as prognostic value is not assessed in this work (Section 4.1).

All five are strongly differentially expressed on raw read counts, and the per-sample values in Appendix B show the tumor and normal distributions to be largely non-overlapping rather than separated by a few outlying samples. The support is uneven, however: *hsa-mir-888* is detected in only one of five normal samples, so its upregulation rests on near-absence in a small baseline, and *hsa-mir-1258* is near zero in half the tumor samples. Each has at most one prior glioma report under an alias-expanded literature search, and none is indexed in the curated biomarker set. That evidence is independent of the model: it rests on the differential-expression contrast and the literature screen, neither of which uses the node-type structure examined in Section 3.2. What the controls do bear on is the ordering. Because the model discriminates only weakly within the miRNA subgraph, the rank and confidence columns of Table 4 should be read as descriptive rather than as an independent line of evidence, and we make no claim that these five are better candidates than other differentially expressed miRNA loci the screen also retained. Prognostic value is separate again and is not assessed here; see Section 4.1.

## 4 DISCUSSION

Glioblastoma is characterized by high mortality and marked tumor heterogeneity, and has few established biomarkers, making tailored diagnosis and treatment difficult. ceRNA analysis offers a way to study the gene regulatory system underlying glioblastoma, but current methods largely depend on statistical measures that assume linear relationships and can therefore misrepresent gene interactions.

To address this gap, we modeled miRNA-lncRNA-mRNA interactions as a heterogeneous graph and trained an RGCN on it to identify high-confidence biomarkers for prognosis and diagnosis. Among the architectures trained on the ceRNA graph, the RGCN discriminated best, and its margin over the convolutional and attention-based alternatives is consistent with the relational structure carrying information that relation-agnostic operators discard. The strongest non-graph baselines, however, were competitive: the hierarchical neural network and random forest reached AUCROC values within the RGCN’s fold-to-fold variation. So the evidence supports the RGCN as the best of the graph architectures tested, but not as categorically superior to all conventional methods. Zhi et al. (2025) likewise found RGCNs effective for biomarker identification via ceRNA analysis, though in gastric cancer; to our knowledge this is the first application of RGCNs to ceRNA network analysis in glioblastoma.

The controls of Section 3.2 change how both results should be read. We had interpreted the fusion ordering — late fusion improving on the ceRNA-only RGCN while early and intermediate fusion did not — as evidence that the two data types are informative about different nodes and are best combined at the decision level. The controls do not support that reading. Copy number is undefined at all 314 miRNA loci, and 27 of the 31 positives sit on nodes with no copy-number measurement, so the `cnv_observed` flag functions as a proxy for the node type that carries almost all of the positive labels. That flag alone reaches an AUCROC of 0.697, reproducing the CNV-only result to within 0.001, while the copy-number value with the flag removed reaches 0.475 *±* 0.007. The CNV branch was therefore contributing observability, not genomic dosage, and the fusion gain is most economically explained as the node-type prior re-entering the ensemble through that channel rather than as complementary biological signal.

The same prior accounts for most of the headline discrimination. A node-type-only baseline with no topology reaches 0.824 *±* 0.139, and an RGCN retrained on labels permuted within node type matches the observed PR-AUC and MCC. We take this as the central finding of the study: on this graph and this label set, most of the performance that would ordinarily be reported is attributable to which node type the labels sit on. The controls are also sensitive enough to detect what remains. The RGCN exceeds its retrained null on AUCROC (*p* = 0.015), so the graph carries some information beyond node type, but that signal is modest and is not resolved within the miRNA subgraph (*p* = 0.10), where 26 positives give limited power. Separating these two components is the contribution of the controls; neither is visible in the metrics these studies conventionally report.

This does not show that ceRNA topology is uninformative in general. It shows that, without these controls, an evaluation of this kind cannot separate topology from label composition, and that the design admits a confound that would inflate any model in this class. We have not tested other datasets, but any study whose positive labels are concentrated in one node type can check for the same effect by reporting a node-type-only baseline, a within-type evaluation, and a retrained within-type permutation null. A fair evaluation would need labels distributed across node types, or would need to be conducted within a single type, and would need a covariate for measurement availability wherever a modality is systematically missing from one type.

### 4.1 Limitations, Future Directions, and Implications

#### Prognostic validation

We do not report a prognostic assessment of the five candidate loci. The only survival analysis available to us, on an independent cohort with matched miRNA expression and survival data, was underpowered for this purpose and null across the miRNAs it covered, so we do not present it as evidence either for or against the candidates’ clinical relevance. Establishing prognostic value will require a larger, adequately powered cohort with matched expression and outcome data.

#### Label set provenance and versioning

The 143-locus curated set used for labeling was assembled from the Cancer Gene Census, GeneCards, DisGeNET, and the glioblastoma literature, and comprises 67 mRNA, 48 miRNA, and 28 lncRNA entries, of which 31 map onto graph nodes. The compiled set is released with this work, but it was assembled without a recorded per-entry citation or a pinned database version, so it cannot be regenerated exactly from its sources as it stands. Its miRNA enrichment is the proximate cause of the confound in Section 3.2, which makes its provenance load-bearing rather than incidental; a versioned, per-entry-sourced replacement is the single change most likely to alter the conclusions here.

#### Novelty screen naming coverage

The screen expands each locus to its conventional mature-strand, bare-numeric and copy-independent names, which recovers records a precursor-only search misses; the three well-characterized loci rise from 3, 1 and 0 records to 19, 12 and 11 under expansion. It does not, however, consult the full miRBase alias table, so a locus discussed exclusively under a historical or withdrawn identifier could still be missed. The reported counts remain lower bounds, though the two candidates whose counts changed under expansion suggest the residual gap is small.

#### Capacity asymmetry in the fusion comparison

The late-fusion arm is not capacity-matched to the ceRNA-only baseline. Its graph branch is the identical RGCN, but it adds an independently trained random forest of 300 trees and a per-fold mixing weight, so the comparison confounds decision-level fusion with the addition of a second model and with ensembling. An ensemble of two RGCNs would be the appropriate control and was not run.

#### Cohort composition and IDH status

The analysis was run over TCGA-GBM without stratifying on IDH mutation status. The Xena TCGA-GBM clinical matrix carries no direct IDH mutation call, so we report the glioma CpG island methylator phenotype instead, which is closely associated with IDH mutation: of 629 samples, 505 are non-G-CIMP, 47 are G-CIMP, and 77 are unlabeled. Taking G-CIMP as a proxy, on the order of 7% of the labeled cohort is likely IDH-mutant. Under the 2021 WHO classification those tumors would now be graded as astrocytoma rather than glioblastoma, so the cohort is not a homogeneous IDH-wildtype glioblastoma population by current criteria, although the contamination is modest. G-CIMP status enters the model only as one of the nine clinical node-feature axes (Appendix A) and was not used to filter samples. Restricting to IDH-wildtype cases, using a direct mutation call rather than a methylation proxy, and repeating the differential-expression step would be required before the gene set is described as glioblastoma-specific under current criteria.

#### Label imbalance and node-type confounding

Of 143 curated glioblastoma biomarkers, only 31 map onto nodes in the graph, a positive rate of 1.49%. The positives are unevenly distributed across node types: 26 of the 31 are miRNA, so 8.28% of miRNA nodes are labeled positive against 0.34% of lncRNA and 0.26% of mRNA nodes. A miRNA node is therefore roughly 32 times more likely to carry a positive label than an mRNA node. Node type is visible to the model through the relation structure. Section 3.2 quantifies the consequence: a type-only baseline recovers most of the reported AUCROC, the retrained permutation null matches the observed PR-AUC, and within the miRNA subgraph discrimination is weak. This is no longer a caveat on the headline number but the main explanation for it. The limitation is a property of the label set and graph construction rather than of the architecture, and it would apply to any model in this class evaluated the same way.

#### Cohort and feature constraints

Training data came solely from TCGA-GBM, which largely represents Western populations, so generalizability may be limited. Few clinical features were available, and the lineage plasticity of glioblastoma means that genes identified here may not be viable therapeutic targets. Because GTEx provides no miRNA profiles, only five normal-tissue miRNA samples were available, which limited the statistical power of every miRNA analysis.

#### Graph construction

The graph’s edges are unweighted and do not account for binding affinity or thermodynamic stability, so some modeled interactions may not be biophysically viable.

#### Interpretability

Interpretability is a key challenge in adopting deep learning over statistical methods. Rudin (2019) argues that interpretable models are preferable to black-box models in high-stakes domains. While this argument holds for linear or tabular data, it is less straightforward in molecular biology, because ceRNA networks form an extensive, multilayered regulatory system (Sumazin et al., 2011). RGCNs address this through their message-passing and aggregation mechanism, which condenses many input features into node representations, but the mechanism remains a black box in practice: the RGCN’s limited interpretability obscures which graph features drive predictions. To bridge this gap, future work could employ explainer methods such as GNNExplainer (Ying et al., 2019), which quantifies the influence of input components on the output.

The first question this work leaves open is whether ceRNA topology carries biomarker signal at all under an evaluation that can detect it. Answering it needs a label set that is not concentrated on one node type, or an evaluation conducted entirely within a single type at a sample size that supports it; the miRNA subgraph here offers 26 positives, which is too few to establish a negative. Two further questions follow: how far the results depend on the permissive co-regulation rule used to infer two thirds of the edges, which an edge-removal ablation would settle, and whether a modality missing systematically from one node type can be included without reintroducing the observability confound. Beyond that, an inductive evaluation on nodes held out of the graph entirely would test generalization in a way the present transductive protocol cannot. We would encourage the controls reported here to be run as a matter of course in ceRNA graph-learning studies, since they are inexpensive and, on this dataset, changed the interpretation of every headline result.

## 5 CONCLUSION

We applied relational graph convolutional networks to a heterogeneous ceRNA network in glioblastoma, integrated with gene-level copy-number data through three fusion strategies, and then tested whether the resulting performance reflected ceRNA topology. For the most part it does not. A node-type-only baseline using no topology reaches an AUCROC of 0.824 *±* 0.139; an RGCN retrained on labels permuted within node type matches the observed PR-AUC (*p* = 0.33), and exceeds the null only modestly on AUCROC (*p* = 0.015); and within the miRNA subgraph, where the node-type prior is constant, discrimination is weak. The apparent benefit of decision-level CNV fusion has the same origin: copy number is undefined at every miRNA locus, the observability flag alone reaches 0.697, and the copy-number value with that flag removed reaches 0.475 *±* 0.007.

The contribution is methodological. We provide a reproducible heterogeneous-graph framework for ceRNA-based prioritization — graph construction, labels, fusion variants, and the controls above — and, with it, a set of controls that separates network signal from label composition. Curated biomarker sets are often enriched for one molecular species, and heterogeneous graphs encode node type in their relation structure; wherever both hold, the same check applies. The controls are inexpensive, need no data beyond what the study already has, and here substantially changed the conclusion the uncontrolled metrics would have supported. We would encourage them as routine practice in this literature, and we release everything needed to run them.

The five candidate loci of Section 3.3 stand on their differential expression, which holds when re-estimated from raw read counts, and on the literature screen, both of which are independent of the model and of the confound; the evidence for *hsa-mir-888* and *hsa-mir-1258* is weaker than for the other three. What we do not claim is that the model’s ranking adds evidence beyond it, or that the loci are prognostic; establishing the latter requires the matched expression and outcome data discussed in Section 4.1.

## ETHICS STATEMENT

This study analyzed only de-identified, publicly available human data obtained from The Cancer Genome Atlas and the Genotype-Tissue Expression project. Both resources obtained informed consent and ethical approval at the contributing institutions, and all data were released in anonymized form in accordance with their respective data-use policies. No new human or animal data were generated, and no identifiable participant information was accessed, so additional ethical approval and written informed consent were not required for this work.

## AUTHOR CONTRIBUTIONS

SK: Conceptualization, Data curation, Formal Analysis, Investigation, Methodology, Project administration, Resources, Software, Validation, Visualization, Writing – original draft, Writing – review and editing. JZ: Funding acquisition, Supervision, Writing – review and editing. NJ: Supervision, Writing – review and editing. All authors contributed to the article and approved the submitted version.

## DATA AVAILABILITY STATEMENT

This study analyzed publicly available datasets. Tumor (TCGA-GBM primary tumor) and normal (GTEx, all brain regions) mRNA and lncRNA expression, as RSEM expected counts, were obtained from the UCSC Xena Toil recompute hub (https://toil.xenahubs.net; datasets `TcgaTargetGtex_gene_expected_count` and `TcgaTargetGTEX_phenotype.txt`). TCGA-GBM miRNA-seq read counts were obtained from the NCI Genomic Data Commons (data type “miRNA Expression Quantification”; https://portal.gdc.cancer.gov/projects/TCGA-GBM); the graph’s original miRNA contrast used the UCSC Xena GDC hub matrix (https://gdc.xenahubs.net; dataset `TCGA-GBM.mirna.tsv`). GISTIC2 thresholded gene-level copy-number calls and clinical data were obtained from the UCSC Xena TCGA hub (https://tcga.xenahubs.net; datasets TCGA.GBM.sampleMap/Gistic2 CopyNumber Gistic2 all thresholded.by genes and TCGA.GBM.sampleMap/GBM clinicalMatrix). All Xena datasets can be browsed at https://xenabrowser.net (Goldman et al., 2020). The underlying data originate from the NCI Genomic Data Commons (https://portal.gdc.cancer.gov/projects/TCGA-GBM; dbGaP phs000178) and the GTEx Portal (https://gtexportal.org; dbGaP phs000424). Interaction edges were drawn from miRTarBase (Release 9.0; https://mirtarbase.cuhk.edu.cn), LncACTdb (version 3.0; http://bio-bigdata.hrbmu.edu.cn/LncACTdb), and TargetScan (release 8.0; https://www.targetscan.org). ENCORI (https://rnasysu.com/encori/) was queried and contributed no edges. Gene identifiers were mapped with the HGNC complete set (https://www.genenames.org) and MyGene.info API v3 (https://mygene.info). The positive-label set was compiled from the Cancer Gene Census (https://cancer.sanger.ac.uk/census), GeneCards (https://www.genecards.org), DisGeNET (https://www.disgenet.org), and the glioblastoma literature.

All analysis code is publicly available at https://github.com/SamarthKhandelwal-create/RGCN-GBM-Biomarker-Detection, including the graph construction, the curated label set, the fusion variants, and the control experiments of Section 3.2. A versioned archive of the graph, the curated label set, the copy-number vector, the differential-expression output, and the control experiments of Section 3.2 is deposited on Zenodo at https://doi.org/10.5281/zenodo.22599653, which resolves to the most recent version. The archive documents its own limitations, including the label set’s missing per-entry citations and unpinned source versions.

## A CLINICAL NODE FEATURE STRATIFICATION

Each node in the heterogeneous graph carries a vector of binary clinical features derived from the TCGA-GBM metadata. Table A1 lists each clinical variable, the criterion used to stratify patients into two groups, and the rationale for its inclusion.

**Table A1.**
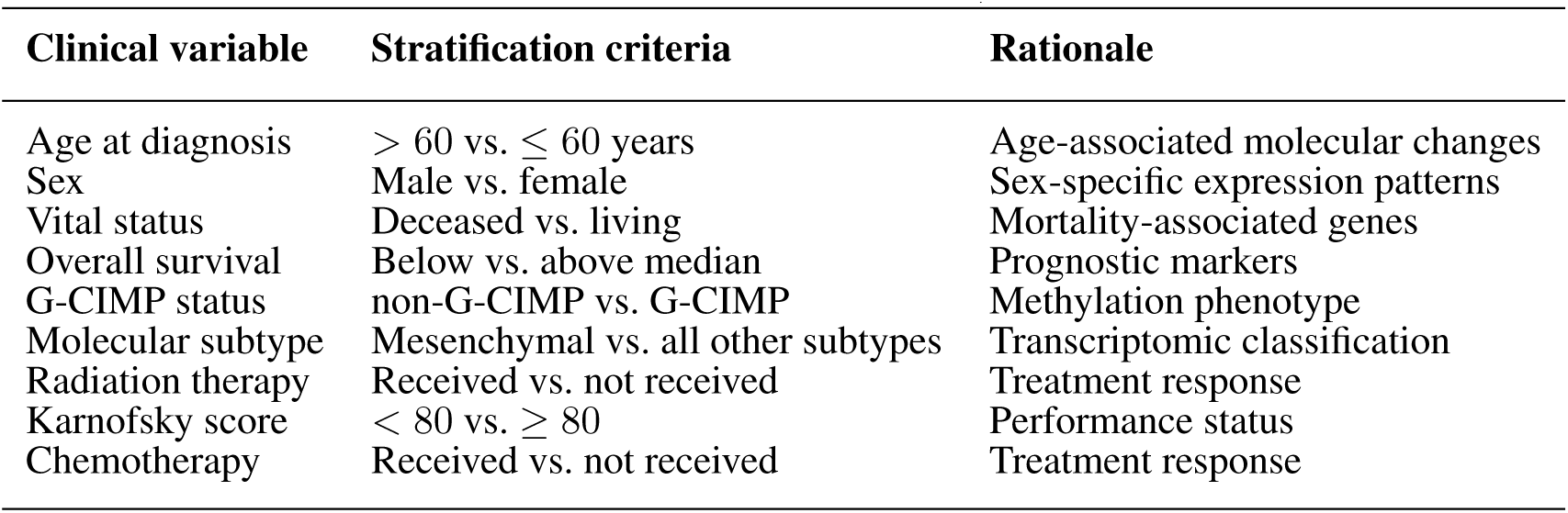
Clinical variables used as node features, with stratification criteria.

| Clinical variable | Stratification criteria | Rationale |
| --- | --- | --- |
| Age at diagnosis | > 60 vs. ≤ 60 years | Age-associated molecular changes |
| Sex | Male vs. female | Sex-specific expression patterns |
| Vital status | Deceased vs. living | Mortality-associated genes |
| Overall survival | Below vs. above median | Prognostic markers |
| G-CIMP status | non-G-CIMP vs. G-CIMP | Methylation phenotype |
| Molecular subtype | Mesenchymal vs. all other subtypes | Transcriptomic classification |
| Radiation therapy | Received vs. not received | Treatment response |
| Karnofsky score | < 80 vs. ≥ 80 | Performance status |
| Chemotherapy | Received vs. not received | Treatment response |

Two of these criteria warrant clarification. The methylation variable is the TCGA `G CIMP STATUS` field, which records the glioma CpG island methylator phenotype; although G-CIMP is closely associated with IDH1/2 mutation, the two are distinct measurements and are reported separately by TCGA. For molecular subtype (Verhaak et al., 2010), the Mesenchymal group is contrasted against all other assigned subtypes (Classical, Neural, and Proneural), not against Proneural alone. Each stratification is oriented so that group 1 denotes the higher-risk or more aggressive condition, which is why the Karnofsky criterion inverts the direction of its numeric threshold.

## B PER-SAMPLE EXPRESSION OF THE CANDIDATE LOCI

The miRNA differential-expression contrast rests on 6 tumor and 5 normal samples (Section 2.1.1). Because that is a small design and the fold changes reported in Table 4 are large, Table A2 gives the underlying per-sample values so the reader can judge the evidence directly rather than through the summary statistics.

Values are log_2_(CPM + 1) computed from the GDC raw read counts used for the differential-expression analysis.

**Table A2.** Per-sample expression of the five candidate loci, in log_2_(CPM + 1). Detection is the number of samples with a non-zero value.

| Locus | Tumor median<br>(n = 6) | Normal median<br>(n = 5) | Detected, tumor | Detected, normal |
| --- | --- | --- | --- | --- |
| hsa-mir-1258 | 0.03 | 3.95 | 3/6 | 5/5 |
| hsa-mir-203b | 0.33 | 5.16 | 6/6 | 5/5 |
| hsa-mir-450a-2 | 7.25 | 1.59 | 6/6 | 5/5 |
| hsa-mir-888 | 3.59 | 0.00 | 6/6 | 1/5 |
| hsa-mir-5683 | 0.36 | 4.49 | 5/6 | 5/5 |

The direction of every locus is consistent across samples, and the tumor and normal distributions are largely non-overlapping, which is why the fold changes are large. Two cautions follow from the table rather than from the summary statistics. First, *hsa-mir-888* is detected in only one of five normal samples, so its apparent upregulation rests on near-absence in a five-sample baseline rather than on a well-estimated normal level. Second, *hsa-mir-1258* is detected in half the tumor samples at values near zero, so its downregulation is driven by the normal samples being uniformly positive rather than by a graded difference.

## C BIOLOGICAL CONTEXT FOR THE CANDIDATE LOCI

None of the five loci has an established role in glioblastoma; that is why the literature screen retained them. Each does, however, have some characterization in other tissues, which is the most that can be said about them at present. This appendix summarizes that context so the candidates can be prioritized for follow-up. Given Section 3.2, we present it as background for loci that are differentially expressed in this cohort, not as support for a model-derived prediction.

### hsa-mir-1258 (downregulated, log_2_FC −5.64)

miR-1258 has been reported as a suppressor of invasion and metastasis in several epithelial cancers, and is described as acting through the miR-1258/RUNX1 axis in a circular-RNA sponging context, characterized in rheumatoid arthritis synovial fibroblasts rather than in a tumor (Xu et al., 2023). A tumor-suppressive role is consistent with the direction observed here, and the sponging context is the kind of interaction the ceRNA framework is meant to capture, which makes this locus the most mechanistically coherent of the five. Its low assay abundance (base mean 56.8, the lowest of the five) is the main caveat.

### hsa-mir-203b (downregulated, log_2_FC −6.03)

The miR-203 family is a well-studied epithelial differentiation regulator, but the *203b* paralogue specifically is sparsely characterized, and it has recently appeared in miRNA signature panels for non-small-cell lung cancer diagnosis (Zhang et al., 2026). It has the cleanest per-sample separation of the five (Table A2) and no prior glioma report at all, which makes it the strongest candidate for experimental follow-up on expression grounds.

### hsa-mir-450a-2 (upregulated, log_2_FC +6.04)

miR-450a has been reported in gastric cancer, including in a machine-learning liquid-biopsy panel predicting chemotherapy response (Shoda et al., 2025). The -2 suffix denotes one of two genomic loci producing the same mature product, so any functional claim would need to establish which locus is transcribed. It has the highest abundance of the five (base mean 847.8) and is detected in every sample, so its effect estimate is the best supported.

### hsa-mir-888 (upregulated, log_2_FC +7.55)

miR-888 is the founding member of a primate-specific cluster characterized as an oncogenic network in prostate (Hasegawa et al., 2018), where elevated expression associates with aggressive disease. An oncogenic role is consistent with the direction here. This locus carries the weakest statistical support of the five: low abundance (base mean 65.3), the largest standard error (1.359), and detection in only one of five normal samples.

### hsa-mir-5683 (downregulated, log_2_FC −6.94)

miR-5683 is essentially uncharacterized. It appears mainly as a component of competing-endogenous-RNA axes identified by bioinformatic screens, for example in gastric cancer (Aslani et al., 2026), rather than in any dedicated functional study. It has the largest absolute fold change of the five and no prior glioma literature, but there is no independent biological evidence to weigh, so it should be regarded as the most speculative candidate.

## D IMBALANCE-AWARE TRAINING

Figure A1 compares precision–recall curves for the reference RGCN, the selected focal-loss and positive-unlabeled variants, and the node-type-only model; Figure A2 reports the threshold-dependent metrics, with thresholds chosen by validation MCC, and the precision among the top-ranked nodes of each test fold. All variants share the folds, seed, and evaluation protocol of Section 2.5.

**Figure A1.**
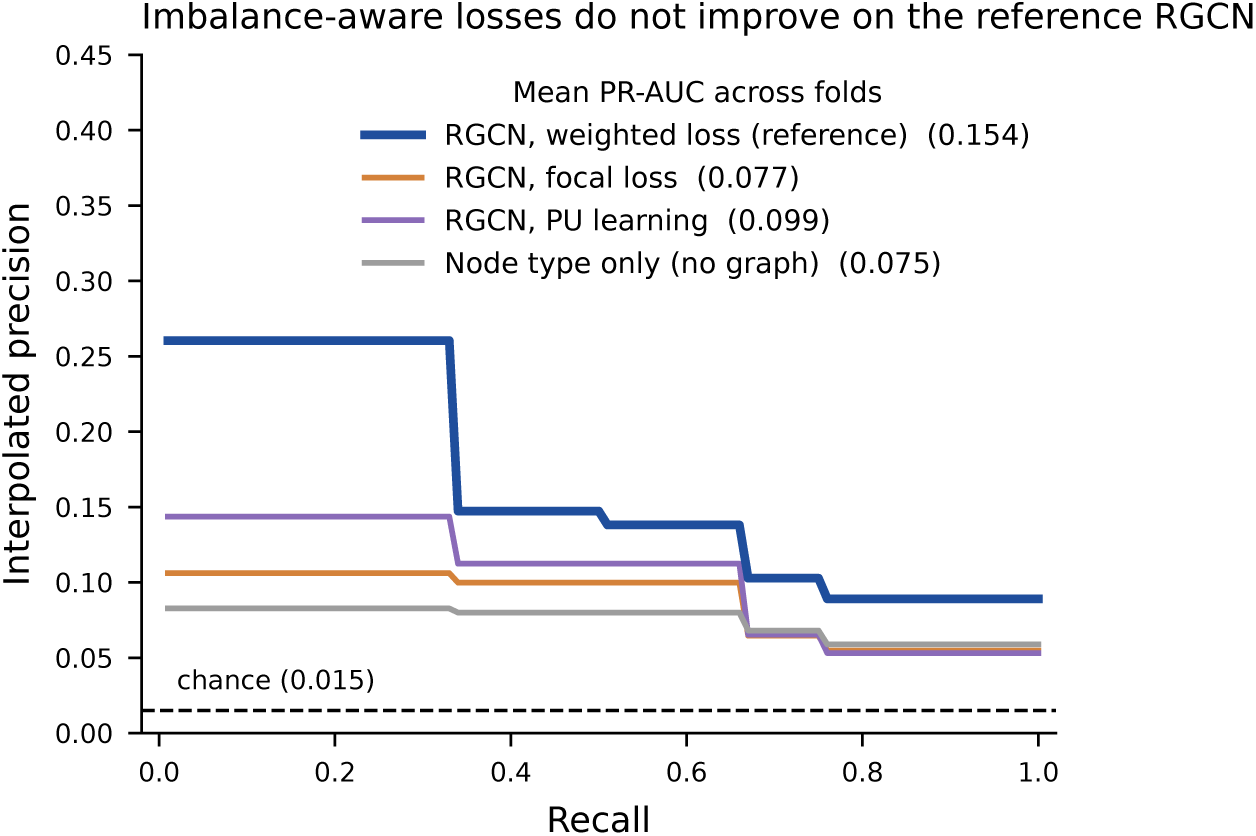
Precision–recall curves (precision interpolated and averaged over the ten test folds) for the reference RGCN, the imbalance-aware variants, and the node-type-only model. The legend gives the mean PR-AUC across folds; the dashed line marks the chance level of 0.015.

## E FUNCTIONAL ENRICHMENT ANALYSIS

Functional enrichment analysis was used to confirm that the differentially expressed genes were biologically coherent rather than statistical artifacts. The recovered terms and pathways are consistent with those reported in prior glioblastoma ceRNA studies (Bazrgar et al., 2024; Liu et al., 2021), supporting the validity of the processed dataset. Gene Ontology biological process terms are shown in Figure A3, and the corresponding KEGG pathway enrichment in Figure A4.

**Figure A2.**
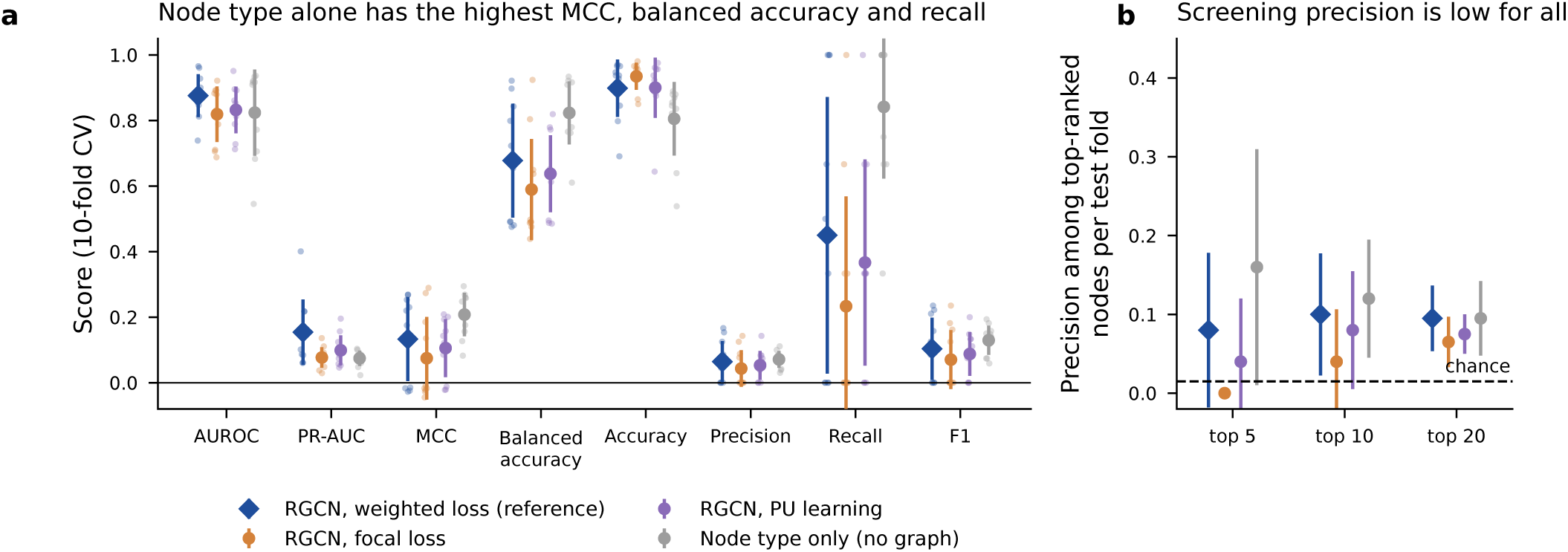
(a) Ranking and threshold-dependent metrics over the ten folds (points are folds; markers and bars are mean *±* SD), with thresholds selected by validation MCC. (b) Precision among the top 5, 10 and 20 ranked nodes of each test fold; the dashed line marks the chance level.

**Figure A3.**
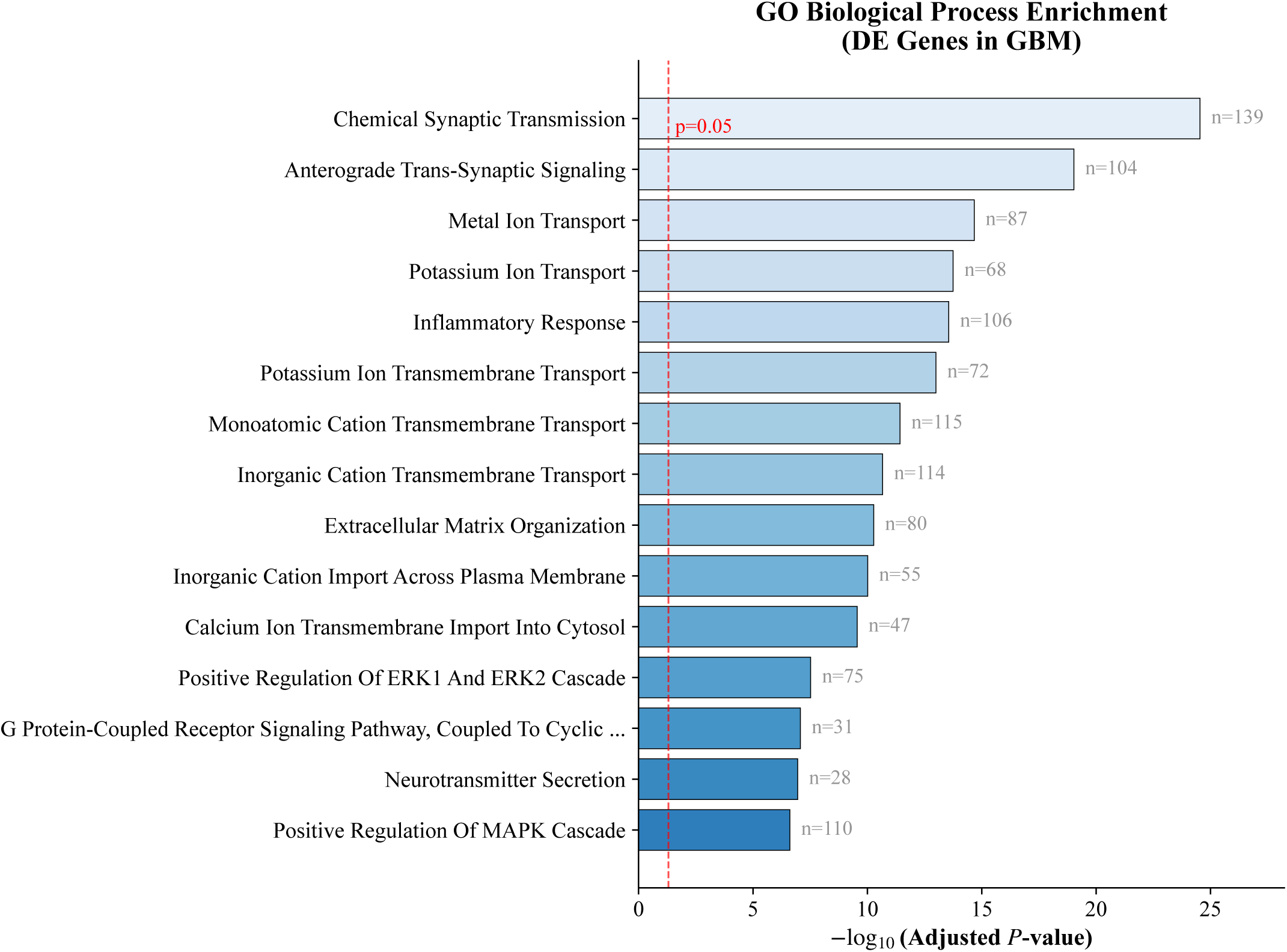
Gene Ontology biological process enrichment for the differentially expressed genes in GBM. Bars show *−* log_10_(adjusted *p*-value); the dashed line marks *p* = 0.05 and *n* denotes the number of genes in each term.

**Figure A4.**
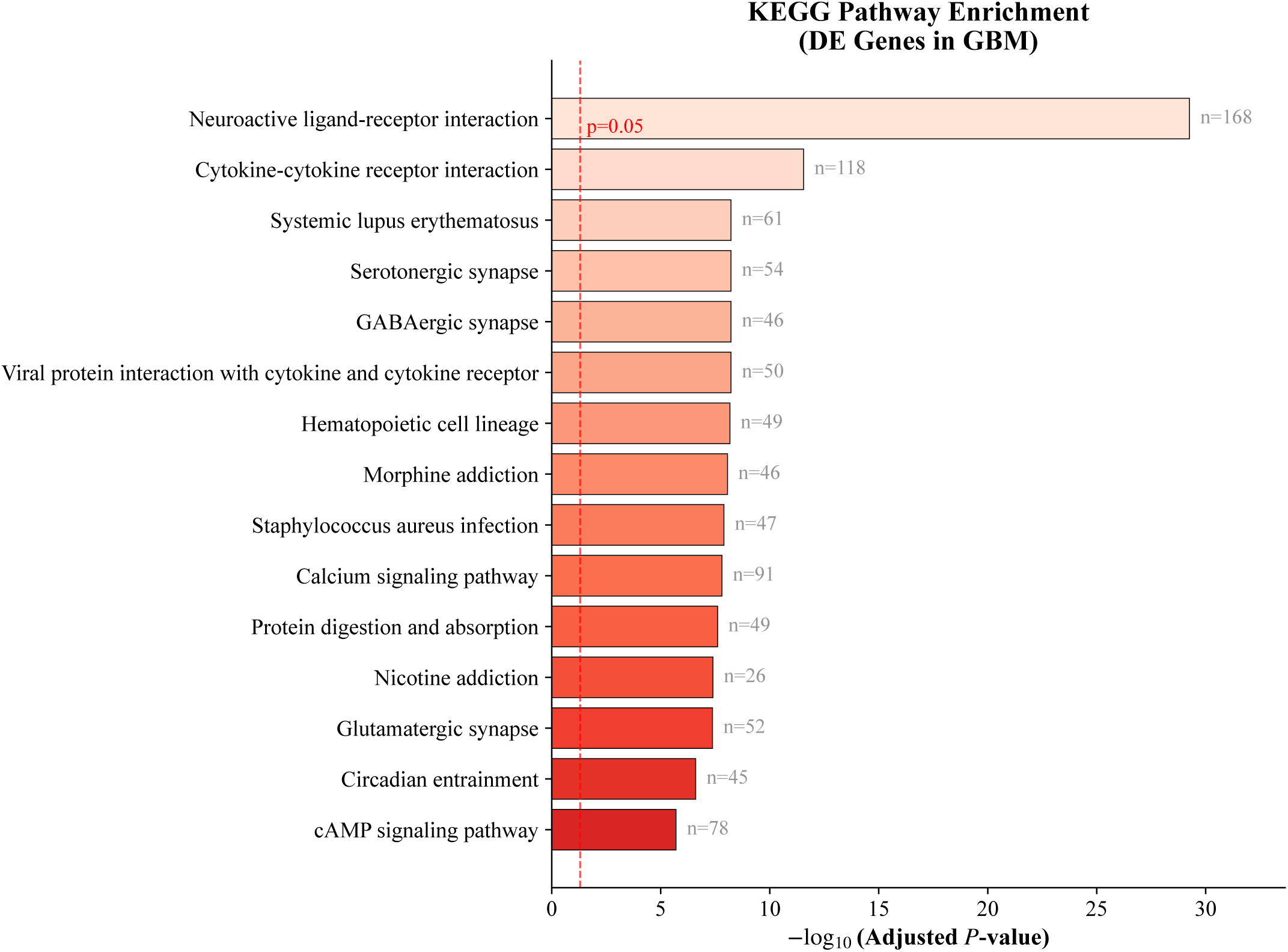
KEGG pathway enrichment for the differentially expressed genes in GBM. Bars show *—* log_10_(adjusted *p*-value); the dashed line marks *p* = 0.05 and *n* denotes the number of genes in each pathway.

## REFERENCES

Agarwal, V., Bell, G. W., Nam, J.-W., and Bartel, D. P. (2015). Predicting effective microRNA target sites in mammalian mRNAs. eLife 4, e05005. doi:10.7554/eLife.05005

Aslani, S., Kalantary-Charvadeh, A., Morovat, P., Abbasalipourkabir, R., Nourmohammadi, I., Emami Razavi, A., et al. (2026). Integrated bioinformatics and tissue-based validation reveal the oncogenic role of hsa circ 0043256 and hsa circ 0004789 in gastric cancer. Biochemistry and Biophysics Reports 45, 102442. doi:10.1016/j.bbrep.2026.102442

Bazrgar, M., Mirmotalebisohi, S. A., Ahmadi, M., Azimi, P., Dargahi, L., Zali, H., et al. (2024). Comprehensive analysis of lncRNA-associated ceRNA network reveals novel potential prognostic regulatory axes in glioblastoma multiforme. Journal of Cellular and Molecular Medicine 28, e18392. doi:10.1111/jcmm.18392

Bray, F., Laversanne, M., Sung, H., Ferlay, J., Siegel, R. L., Soerjomataram, I., et al. (2024). Global cancer statistics 2022: GLOBOCAN estimates of incidence and mortality worldwide for 36 cancers in 185 countries. CA: A Cancer Journal for Clinicians 74, 229–263. doi:10.3322/caac.21834

Chicco, D. and Jurman, G. (2020). The advantages of the Matthews correlation coefficient (MCC) over F1 score and accuracy in binary classification evaluation. BMC Genomics 21, 6. doi:10.1186/s12864-019-6413-7

Goldman, M. J., Craft, B., Hastie, M., Repecka, K., McDade, F., Kamath, A., et al. (2020). Visualizing and interpreting cancer genomics data via the Xena platform. Nature Biotechnology 38, 675–678. doi:10.1038/s41587-020-0546-8

Hasegawa, T., Glavich, G. J., Pahuski, M., Short, A., Semmes, O. J., Yang, L., et al. (2018). Characterization and evidence of the miR-888 cluster as a novel cancer network in prostate. Molecular Cancer Research 16, 669–681. doi:10.1158/1541-7786.MCR-17-0321

Huang, H.-Y., Lin, Y.-C.-D., Li, J., Huang, K.-Y., Shrestha, S., Hong, H.-C., et al. (2022). miRTarBase update 2022: An informative resource for experimentally validated miRNA-target interactions. Nucleic Acids Research 50, D222–D230. doi:10.1093/nar/gkab1079

Liu, G., Liu, D., Huang, J., Li, J., Wang, C., Liu, G., et al. (2021). Comprehensive analysis of ceRNA network related to lincRNA in glioblastoma and prediction of clinical prognosis. BMC Cancer 21, 98. doi:10.1186/s12885-021-07817-5

Mermel, C. H., Schumacher, S. E., Hill, B., Meyerson, M. L., Beroukhim, R., and Getz, G. (2011). GISTIC2.0 facilitates sensitive and confident localization of the targets of focal somatic copy-number alteration in human cancers. Genome Biology 12, R41. doi:10.1186/gb-2011-12-4-r41

Muzellec, B., Telenczuk, M., Cabeli, V., and Andreux, M. (2023). PyDESeq2: A Python package for bulk RNA-seq differential expression analysis. Bioinformatics 39, btad547. doi:10.1093/bioinformatics/btad547

Neftel, C., Laffy, J., Filbin, M. G., Hara, T., Shore, M. E., Rahme, G. J., et al. (2019). An integrative model of cellular states, plasticity, and genetics for glioblastoma. Cell 178, 835–849.e21. doi:10.1016/j.cell.2019.06.024

Ostrom, Q. T., Price, M., Neff, C., Cioffi, G., Waite, K. A., Kruchko, C., et al. (2023). CBTRUS statistical report: Primary brain and other central nervous system tumors diagnosed in the united states in 2016–2020. Neuro-Oncology 25, iv1–iv99. doi:10.1093/neuonc/noad149

Roncevic, A., Koruga, N., Koruga, A. S., and Roncevic, R. (2025). Why do glioblastoma treatments fail? Future Pharmacology 5, 7. doi:10.3390/futurepharmacol5010007

Rudin, C. (2019). Stop explaining black box machine learning models for high stakes decisions and use interpretable models instead. Nature Machine Intelligence 1, 206–215. doi:10.1038/s42256-019-0048-x

Saito, T. and Rehmsmeier, M. (2015). The precision-recall plot is more informative than the ROC plot when evaluating binary classifiers on imbalanced datasets. PLOS ONE 10, e0118432. doi:10.1371/journal.pone.0118432

Salmena, L., Poliseno, L., Tay, Y., Kats, L., and Pandolfi, P. P. (2011). A cerna hypothesis: The rosetta stone of a hidden rna language? Cell 146, 353–358. doi:10.1016/j.cell.2011.07.014

Schlichtkrull, M., Kipf, T. N., Bloem, P., van den Berg, R., Titov, I., and Welling, M. (2017). Modeling relational data with graph convolutional networks. arXiv preprint arXiv:1703.06103

Shoda, K., Xu, C., Nagasaka, T., Ichikawa, D., and Goel, A. (2025). A machine-learning powered liquid biopsy predicts response to paclitaxel plus ramucirumab in advanced gastric cancer: Results from the prospective IVY trial. Molecular Cancer 25, 30. doi:10.1186/s12943-025-02526-0

Su, Y., Liang, C., and Yang, Q. (2021). LncRNA MALAT1 promotes glioma cell growth through sponge miR-613. Journal of B.U.ON. 26, 984–991

Sumazin, P., Yang, X., Chiu, H.-S., Chung, W.-J., Iyer, A., Llobet-Navas, D., et al. (2011). An extensive microRNA-mediated network of RNA-RNA interactions regulates established oncogenic pathways in glioblastoma. Cell 147, 370–381. doi:10.1016/j.cell.2011.09.041

The Cancer Genome Atlas Research Network (2008). Comprehensive genomic characterization defines human glioblastoma genes and core pathways. Nature 455, 1061–1068. doi:10.1038/nature07385

The GTEx Consortium (2013). The genotype-tissue expression (GTEx) project. Nature Genetics 45, 580–585. doi:10.1038/ng.2653

Verhaak, R. G. W., Hoadley, K. A., Purdom, E., Wang, V., Qi, Y., Wilkerson, M. D., et al. (2010). Integrated genomic analysis identifies clinically relevant subtypes of glioblastoma characterized by abnormalities in PDGFRA, IDH1, EGFR, and NF1. Cancer Cell 17, 98–110. doi:10.1016/j.ccr.2009.12.020

Wang, P., Guo, Q., Qi, Y., Hao, Y., Gao, Y., Zhi, H., et al. (2022). LncACTdb 3.0: An updated database of experimentally supported ceRNA interactions and personalized networks contributing to precision medicine. Nucleic Acids Research 50, D183–D189. doi:10.1093/nar/gkab1092

Wu, C., MacLeod, I., and Su, A. I. (2013). BioGPS and MyGene.info: Organizing online, gene-centric information. Nucleic Acids Research 41, D561–D565. doi:10.1093/nar/gks1114

Xu, Y., Zai, Z., Lu, Z., Zhang, T., Wang, L., Qian, X., et al. (2023). Circular RNA CircCDKN2B-AS 006 promotes the tumor-like growth and metastasis of rheumatoid arthritis synovial fibroblasts by targeting the miR-1258/RUNX1 axis. International Journal of Molecular Sciences 24, 5880. doi:10.3390/ijms24065880

Ying, Z., Bourgeois, D., You, J., Zitnik, M., and Leskovec, J. (2019). GNNExplainer: Generating explanations for graph neural networks. In Advances in Neural Information Processing Systems. vol. 32, 9240–9251

Zhang, Y., Wang, Y., Ma, D., Xiao, G., Feng, L., Cai, J., et al. (2026). Machine learning-advanced hydrogel-based transcription-coupled positive-feedback CRISPR/Cas13a analysis for novel microRNA signatures in differential diagnosis of non-small cell lung cancer. Journal of Nanobiotechnology 24, 557. doi:10.1186/s12951-026-04476-1

Zhi, P., Liu, Y., Zhao, C., and He, K. (2025). GCBRGCN: Integration of ceRNA and RGCN to identify gastric cancer biomarkers. Bioengineering 12, 255. doi:10.3390/bioengineering12030255

